# A shared spatial topography links the functional connectome correlates of cocaine use disorder and dopamine D_2/3_ receptor densities

**DOI:** 10.1101/2023.11.17.567591

**Authors:** Jocelyn A. Ricard, Loïc Labache, Ashlea Segal, Elvisha Dhamala, Carrisa V. Cocuzza, Grant Jones, Sarah Yip, Sidhant Chopra, Avram J. Holmes

## Abstract

**Background:** The biological mechanisms that contribute to cocaine and other substance use disorders involve an array of cortical and subcortical systems. Prior work on the development and maintenance of substance use has largely focused on cortico-striatal circuits, with relatively less attention on alterations within and across large-scale functional brain networks, and associated aspects of the dopamine system. The brain-wide pattern of temporal co-activation between distinct brain regions, referred to as the functional connectome, underpins individual differences in behavior. Critically, the functional connectome correlates of substance use and their specificity to dopamine receptor densities relative to other metabotropic receptors classes remains to be established.

**Methods:** We comprehensively characterized brain-wide differences in functional connectivity across multiple scales, including individual connections, regions, and networks in participants with cocaine use disorder (CUD; n=69) and healthy matched controls (n=62), Further, we studied the relationship between the observed functional connectivity signatures of CUD and the spatial distribution of a broad range of normative neurotransmitter receptor and transporter bindings as assessed through 18 different normative positron emission tomography (PET) maps.

**Results:** Our analyses identified a widespread profile of functional connectivity differences between individuals with CUD and matched healthy comparison participants (8.8% of total edges; 8,185 edges; p_FWE_=0.025). We largely find lower connectivity preferentially linking default network and subcortical regions, and higher within-network connectivity in the default network in participants with CUD. Furthermore, we find consistent and replicable associations between signatures of CUD and normative spatial density of dopamine D_2/3_ receptors.

**Conclusions:** Our analyses revealed a widespread profile of altered connectivity in individuals with CUD that extends across the functional connectome and implicates multiple circuits. This profile is robustly coupled with normative dopamine D_2/3_ receptors densities. Underscoring the translational potential of connectomic approaches for the study of *in vivo* brain functions, CUD- linked aspects of brain function were spatially coupled to disorder relevant neurotransmitter systems.

**Key Points:** *Question:* Are there group differences in whole brain functional connectivity between individuals with and without cocaine use disorder, and to what extent do these connectivity patterns relate to the spatial distribution of dopamine (D_2/3_) receptor densities?

*Findings:* The presence of cocaine use disorder is associated with brain-wide functional connectivity alterations that are spatially coupled to the density of dopamine (D_2/3_) receptors.

*Meaning:* A preferential and replicable link exists between the functional connectome correlates of cocaine use disorder and dopamine receptor densities across the brain.

## Introduction

The study and treatment of substance use disorders represents a complex, multifaceted challenge with far-reaching implications for individuals, their families, and our broader society. In particular, increasing prevalence of cocaine use disorder (CUD) substantially contributes to the rising overdose deaths in the United States (1). A fundamental question facing the field of addiction neuroscience concerns the extent to which substance use behaviors emerge through local patterns of activity or are instantiated across the broader large-scale networks of the human brain. While prior foundational work has established cortico-striatal-thalamic circuit disruption as a fundamental feature of substance use disorders (2), consistent with systems-level models of substance use disorders (3), striatal circuitry is deeply embedded within spatially distributed and functionally linked systems that span the cortical sheet. Whether alterations in functioning are isolated to specific circuits or diffusely distributed throughout large-scale network architecture remains largely unexplored.

Cocaine preferentially targets the dopamine system, and both tonic and phasic dopamine neurotransmission have been shown to play a critical role in the onset and maintenance of substance use pathology (4). Here, for instance, reduced activity within the large-scale networks supporting attention and inhibitory control points to an imbalance between the core dopaminergic circuits that underlie subjective valuation and conditioned responding and those that support “higher-level” executive functioning. Moreover, the neuromodulatory impact of cocaine is not specific to the dopamine system, while primarily blocking the dopamine transporter and inhibiting its reuptake from the synaptic cleft, it also modulates serotonin and norepinephrine transporters (5). However, the extent to which the brain functional correlates of CUD may be coupled to the spatial distribution of dopaminergic processes, relative to other neurotransmitters, remains to be established.

Here, we investigate the relationship between CUD, whole-brain functional connectivity, and neurotransmitter receptor densities. First, we used the network-based statistic (6) to derive whole-brain functional connectivity differences between individuals with CUD and controls. We then examine the association between the identified functional network and the spatial distribution of receptor densities, inferred from positron emission tomography (PET). In doing so, we demonstrate preferential correspondence between regional connectivity alterations related to CUD and the normative topography of dopamine D_2/3_ receptor densities across three independent PET datasets. These data reliably establish that in CUD, extensive and brain-wide alterations in connectivity exist and are closely coupled with dopamine D_2/3_ receptor densities.

## Methods

### Participants

The current study used data from the SUDMEX CUD imaging dataset (7). A total of n=131 individuals (age range: 18-50), including 69 individuals with CUD (85.51% male) and 62 demographically matched healthy comparison participants (79.03% male) were included in the present study. Notably, these data represent a diverse and non-European-centric population in Mexico City, Mexico. Participants with CUD had to have used for at least one year, with current average use of at least three times per week, with periods of continuous abstinence of less than one month during the last year. Additional participant inclusion criteria can be found in **Supplementary Section 1.** Participant behavioral characteristics and demographics can be found in **Table 1**. The reported study analyses procedures were approved by the Yale University Institutional Review Board IRB #1507016245.

**Table 1:** Demographic characterization of study sample (n=131).

| Group | Cocaine Use Disorder (CUD) | Healthy Comparison |  |
| --- | --- | --- | --- |
| Participants (n, % Male) | 69(85.51) | 62(79.03) | $\chi^2$: 0.30<br>p=0.583 |
| Age (mean±sd) | 31.34±7.27 | 30.42±8.18 | t: -0.68;<br>p=0.501 |
| Education (mean±sd) | 2.83±1.27 | 3.52±1.39 | $\chi^2$: 13.40<br>p<0.020* |
| Head Motion(mean±sd) | 0.23±0.10 | 0.21±0.08 | t: 1.63;<br>p=0.105 |

Three participants were excluded for missing age values. Seven participants were excluded for missing education values. Three participants (2 two HC, 1 CUD) missing sex values, male/female were the only available options. $\chi^2$ , chi-square. Head motion calculated using mean framewise displacement (mm). \* = significant group difference ( $p < 0.05$ ).

### MRI acquisition and processing

Intrinsic (resting state; fcMRI) functional imaging data were acquired using a 3T Phillips Ingenia MR scanner in Mexico City, Mexico. Field-standard processing and quality control procedures were implemented. To generate whole-brain functional connectivity matrices, we parceled each individual’s normalized scans into 400 cortical (8) and 32 subcortical (9) regions. (**Fig. 1A**). Further details can be found in in **Supplement Section 3.**

**Figure 1.**
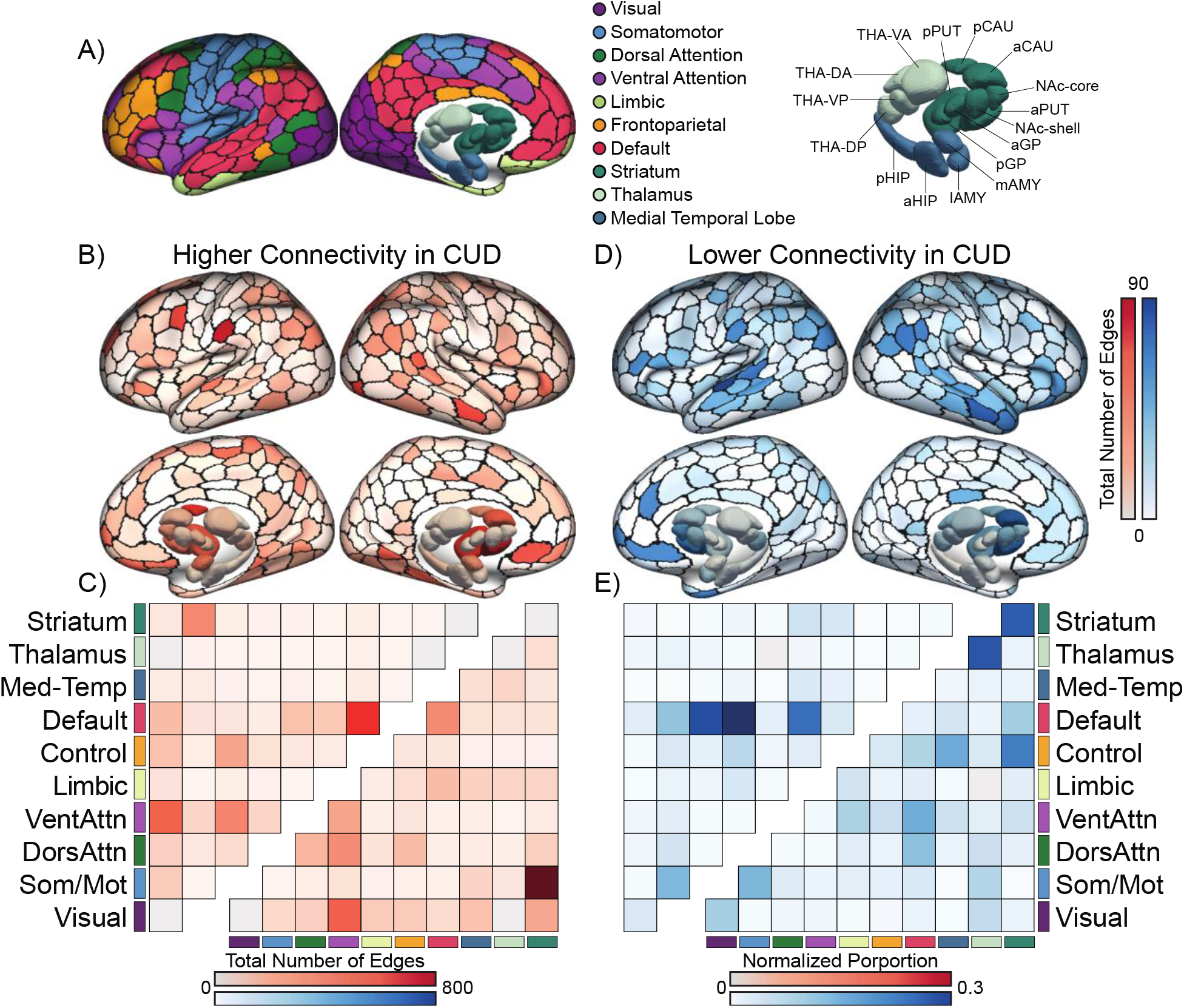
Whole brain atypical functional connectivity in cocaine use disorder (CUD). A widespread network of affected connections exists between individuals with CUD and healthy matched controls, extending across the functional connectome. A) Schaefer 7-network and Tian subcortex parcellations (Scale II) from left to right: a indicates anterior; AMY, amygdala; CAU, caudate nucleus; d, dorsal; DA, dorsoanterior; Default, default network; DorsAttn, dorsal attention network; DP, dorsoposterior; FPN, frontoparietal network; GP, globus pallidus; HIP, hippocampus; l, lateral; Lim, cortical limbic network; m, medial; MTL, medial-temporal lobe (amygdala and hippocampus); NAc, nucleus accumbens; p, posterior; SomMot, somatomotor network; Stri, striatum; PUT, putamen; THA, thalamus. B) Images with a red color scale represent number of significant edges (degree) where individuals with CUD show hyperconnectivity. B) Images with a red color scale represent number of significant negative edges of NBS network where individuals with CUD show hypoconnectivity. C) Heatmap quantified using raw total edge count (upper triangle) and normalized proportion of edges based upon network size (lower triangle) within the NBS component that fall within each of the canonical networks. The darker red indicates higher connectivity in CUD. D) Images with a blue color scale represent number of significant negative edges of NBS network where individuals with CUD show hypoconnectivity. E) Heatmap quantified using raw total edge count (upper triangle) and normalized proportion of edges based upon network size (lower triangle) within the NBS component that fall within each of the canonical networks. Darker blue color indicates lower connectivity in CUD.

### Whole-brain functional connectome correlates of cocaine use disorder

Non-parametric ANCOVA models were used to analyze brain-wide functional connectivity differences between individuals with CUD and matched controls, adjusting for age, sex, and education. The Network Based Statistic (NBS) was used to perform familywise error-corrected (FWE) inference at the level of connected components of edges (12,13), with the primary component-forming threshold, τ, set to *p* < .05 and significance assessed at *p_FWE_* < 0.05. Further statistical details and results for τ = 0.01 and τ = 0.001 are reported in **Supplementary Table 2** and **Supplementary Section 4**.

### Associations between functional dysconnectivity and receptor densities

In order to investigate the relationship between functional alterations identified in individuals with CUD and the topographic distributions of normative neurotransmitter expression in healthy participants, we used Spearman correlation to examined spatial associations between the number of significant connections and normative receptor bindings across each of the 432 - brain regions. These associations were first assessed using 17 unique spatial maps that index a specific receptor or transporter with the largest available sample size (10, 11). Multiple maps from independent datasets were available for some of the receptors and transporters, using either the same or unique tracer. If available, these additional maps were used to assess the stability and replicability of any statistically significant associations (*p* < 0.05). Permutation-based inference (10,000 permutations) using ‘spin-tests’ were used to assess significance, while accounting for spatial autocorrelation. Further statistical details and information regarding specific tracers are provided in **Supplementary Section 1** and **Table 2**.

## Results

### Wide-spread connectivity alterations in cocaine use disorder

We find a significant wide-spread pattern of both hyperconnectivity and hypoconnectivity associated with CUD, encompassing 8.8% of the total edges (8,185 edges; p_FWE_=0.025) linking 432 brain regions (**Fig. 1**). The majority of significant edges (58.94%; 4,824 total edges) demonstrated hypoconnectivity in individuals with CUD. Here, the highest proportion of hypoconnected edges preferentially implicated the default network (**Fig. 1D-E**). After accounting for network size (see **Supplemental Section 4**), connections within striatum and thalamic regions, and between striatum and control networks were preferentially implicated in participants with CUD (**Fig. 1D-E**). At a regional level, precuneus posterior cingulate cortex, medial prefrontal cortex, and anterior caudate nucleus were among the areas most strongly implicated in the network of lower functional connectivity.

When considering patterns of higher connectivity in the CUD group, hyperconnected edges accounted for 41.06% of the total significant edges (3,361 total edges). The total number of hyperconnected edges demonstrated preferential within-network connectivity of the default network, as well as between-network connectivity of striatum and ventral attention networks (**Fig. 1B-C**). When normalizing for the total size of a given network, between-network hyperconnectivity of the striatum-somatomotor networks preferentially emerged. At a regional level, frontal operculum, parietal operculum, extrastriate cortex, and anterior putamen, were among the areas most strongly implicated in the network of higher functional connectivity.

### Shared spatial topography links cocaine use disorder and D_2/3_ receptor densities

Regional functional dysconnectivity was significantly correlated with D_2/3_ receptor density ([^11^C]FLB 457, ρ=0.175 p_spin_=0.015). Associations with D_2/3_ receptors replicated across two additional normative PET maps ([^18^F]fallypride, ρ=0.168; p_spin_=0.022) and ([^11^C]FLB 457, ρ=0.192; p_spin_=0.007) (**Fig. 2B-C**), indicating robust and reliable relationships between D_2/3_ receptors density and CUD-related connectivity dysfunction (**Fig. 2**). To ensure that this association was not driven by large differences in tracer binding between cortical and subcortical regions, we replicated the D_2/3_ receptors association after excluding subcortical regions (**Supplementary Fig. 1**). Associations with two serotonin results were also significant (5HT_4_ [^11^C]SB207145: ρ=0.143; p_spin_=0.032, and 5HT_6_ [^11^C]GSK215083: ρ=0.136; p_spin_=0.020) and reported in **Supplementary Fig. 3,** but did not have replication samples. No associations with other available neurotransmitter systems were detected (**Supplementary Table 2)**.

**Figure 2.**
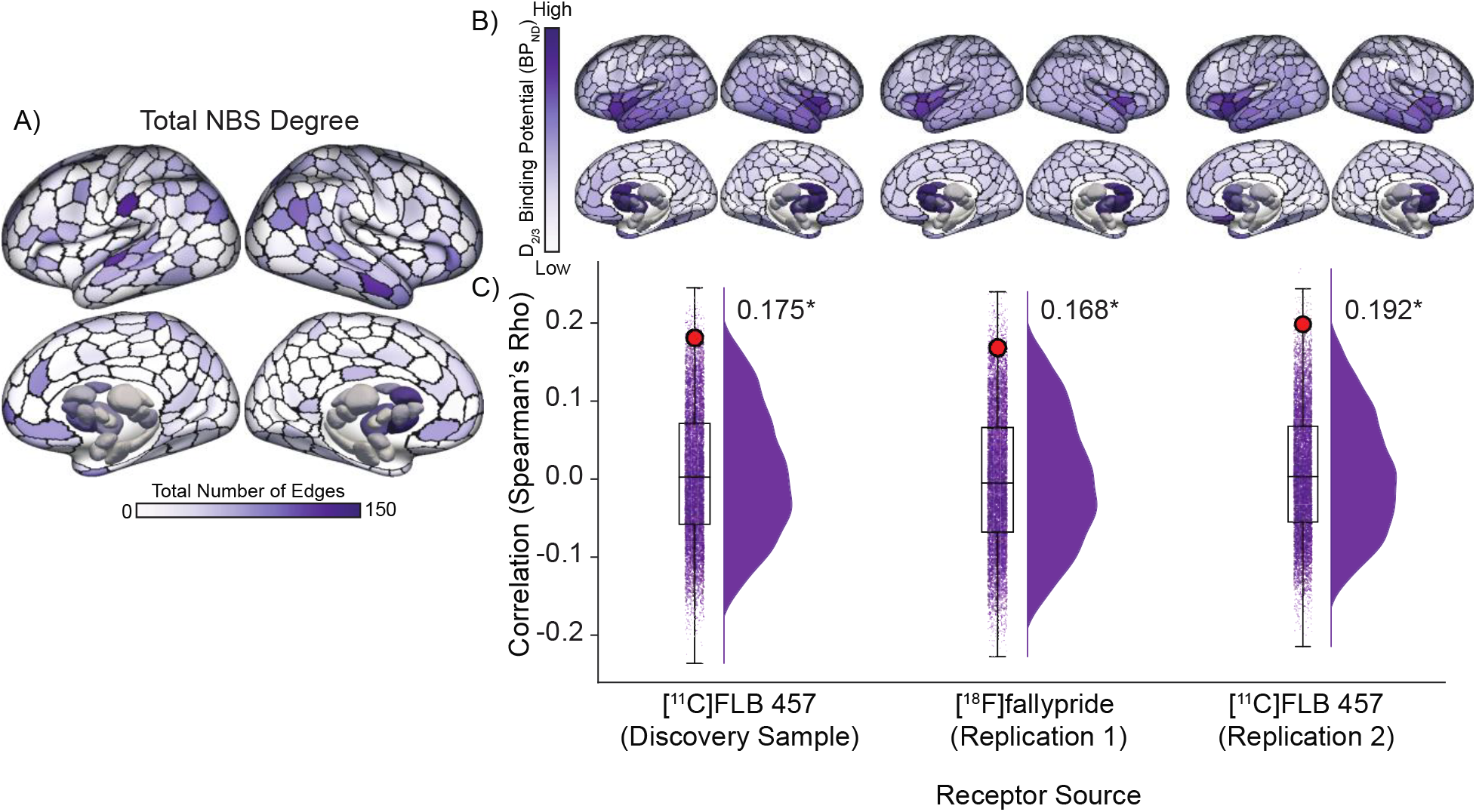
Spatial overlap between whole-brain Network Based Statistic (NBS) network and D_2/3_ receptor density in cocaine use disorder (CUD). A) Visualization of the total (positive and negative) number of significant edges at each region within the NBS component (Fig. 1B + **1D**) where change in fcMRI was significantly correlated with the spatial D_2/3_ receptor density in a discovery sample (Sandiego 2015, p_spin_=0.019) and two replication samples (Jaworska 2020, p_spin_=0.030 and Smith 2017, p_spin_=0.013), respectively). B) D_2/3_ binding potential of PET samples for each receptor source, i.e., discovery sample and replication samples. Color scale normalized between -1.0 to 1.0 for cortex and subcortex separately. C) Each violin-box plot contains (from left to right) distribution of 10k spin-test null correlations between each edge of the NBS component and the spatial density of D_2/3_ receptors. Red dot indicates significant spearman’s correlation. * reflects statistical significance at the threshold p_spin_<0.05. Discovery Sample: Sandiego et al., (2015) (12); Replication 1: Jaworska et al., 2020 (13); Replication 2: Smith et al., 2017 (14).

## Discussion

Cocaine use disorder (CUD) emerges, in part, through the complex interactions of biological systems encompassing neurochemical cascades and associated functional interactions across both local circuits and broader large-scale networks. Establishing how these processes contribute to the onset and maintenance of substance use disorders requires a multi-scale approach, considering measures of *in vivo* brain function, as assessed through fcMRI, as well as neurotransmitter synthesis and transport assessed though PET imaging. In the present analyses, we find wide-spread alterations in intrinsic (“resting-state”) functional connectivity in CUD and by integrating these findings with PET data, we demonstrate the presence of shared spatial patterns linking D_2/3_ receptor densities with the functional connectome correlates of CUD.

While prior work has revealed disruptions in cortico-striatal-thalamic circuitry that underlie varying stages in of substance use disorders (2), our findings support a more diffuse, brain-wide dysregulation in CUD, extending the beyond neural circuit-specific hypotheses. In addition to striatal and thalamic regions, we find alterations in large-scale cortical networks, including the default mode, control, somatomotor, and ventral attention networks, suggesting that dysfunction extends beyond atomically constrained cortico-striatal-thalamic circuitry.

Critically, our findings demonstrate a reliable spatial correspondence between functional dysconnectivity in CUD and the dopaminergic system, extending across both cortical and subcortical regions. Cocaine acts by binding to the dopamine transporter, blocking the reuptake of dopamine from the synaptic cleft, as well as blocking the transporters for norepinephrine and serotonin (5). While our findings also implicate parts of the serotonin system, the most replicable and robust link was found with D_2/3_ receptor densities, suggesting that brain dysconnectivity across large-scale brain networks are preferentially coupled to dopaminergic pathways.

Prior investigations have demonstrated distinguishable functional connectome profiles between CUD and other substance use disorders, such as opioid use disorder, suggesting that individual variability in large-scale connectomes may serve as a valuable predictor for treatment outcomes in CUD (15). While our findings demonstrate a robust pattern of functional alteration in CUD that is coupled with dopaminergic pathways, the extent to which the present findings may reflect a substance specific neurobiological profile or a general profile related to dopaminergic drugs of misuse (for instance, opiates, alcohol, and cocaine) remains to be determined.

The current study, similar to many neuroimaging datasets of individuals with substance use disorders, is limited by its cross-sectional nature and longitudinal approaches may provide further insight on whether neurobiological profiles reflect a vulnerability for illness, a direct consequence of substance use, or the biproduct of illness linked environmental impacts. Moreover, further investigations using concurrent PET and fMRI imaging in patient samples is needed to determine whether illness-related neurochemical alterations interact with brain function. The present sample is also characterized by a large proportion of male participants. Prior work has established the importance of sex differences in the brain-behavior features that characterize substance use disorders (16). Accordingly, data from more sex diverse individuals should be obtained in the future.

## Conclusions

Together, these data establish correspondence across the functional networks implicated in CUD and the neurotransmitters that underlie its mechanism of action. This provides a foundation for future work disentangling the biological mechanisms that govern individual variances in the dopaminergic systems, functional brain organization, and substance use.

## Acknowledgments

This work was supported by: Stanford University Knight-Hennessy Scholars Program (JAR); National Academies of Sciences, Engineering, and Medicine’s Ford Foundation Predoctoral Fellowship (JAR); R01MH120080 (AJH); R01MH123245 (AJH); the Northwell Health Advancing Women in Science and Medicine Career Development Award (ED) and the Feinstein Institutes for Medical Research Emerging Scientist Award (ED); Australian American Association Graduate Fellowship (SC).

## Code Availability

Github: https://github.com/ricardjocelyn/cocaine-use-disorder-receptor-density

## Supplementary Information

### Content

Section 1. Datasets

Section 2. Exclusion Criteria

Section 3. MR Preprocessing

Section 4. Methods (Network Based Statistic, Neuromaps)

Supplementary Table 1: Results using alternate component forming NBS threshold

Supplementary Table 2: Neurotransmitter receptor density spearman’s correlation values

Supplementary Figure 1: Results for 0.01 primary component forming threshold

Supplementary Figure 2: Cortical-only associations and D2/3 spatial density

Supplementary Figure 3: Total NBS degree dysfunction associations with serotonin receptors

Supplementary References

### Section 1. Datasets

We used neuroimaging data from the Mexican magnetic resonance imaging dataset of participants with CUD: SUDMEX CONN (1). SUDMEX is an open-source dataset consisting of 75 (9 female) CUD participants and 62 (11 female) healthy matched controls. Participants were included based on the following inclusion criteria: (a) age between 18 and 50 years old; (b) right-handed; (c) cocaine dependency with an active consumption of at least twice a week in the last month. The exclusion criteria included: (a) current dependence (use and/or abuse) (by DSM-IV criteria) on other substances (alcohol or nicotine); (b) pregnant or breastfeeding; (c) neurological and psychiatric disorders; (d) with a severe systemic disease such as tumors or digestive system disease; and (e) magnetic resonance imaging (MRI) contraindications. All clinical and cognitive assessments were done by trained mental health psychologists and psychiatrists.

### Section 2. Exclusion criteria

Individuals with mean framewise displacement (FD) > 0.55 mm were exclude form analysis. Eliminated 5 participants (one control, four CUD) participants due to artifacts and FD criteria (2).

### Section 3. MR Pre-processing

Briefly, raw images were first put through an automated quality control procedure (3, 4) (fMRIPrep 21.0.1; RRID:SCR_016216), which is based on Nipype 1.6.1 (5, 6) (RRID:SCR_002502). Data were then denoised using aComp-Cor and regressing out six-head motion parameters and mean global signal, followed by high-pass filtering (see below for details). Participants were excluded based on a previously established threshold on framewise displacement (FD; mean FD>0.55mm (2)), visual/manual quality control, and automated MRI quality control pipeline.

#### Preprocessing of B0 inhomogeneity mappings

A B0-nonuniformity map (or field map) was estimated based on two (or more) echo-planar imaging (EPI) references with topup. (7) (FSL 6.0.3:b862cdd5).

#### Anatomical data preprocessing

A total of 1 T1-weighted (T1w) images were found within the input BIDS dataset. The T1- weighted (T1w) image was corrected for intensity non-uniformity (INU) with N4BiasFieldCorrection (8) distributed with ANTs 2.3.3 (9) (RRID:SCR_004757), and used as T1w-reference throughout the workflow. The T1w-reference was then skull-stripped with a Nipype implementation of the antsBrainExtraction.sh workflow (from ANTs), using OASIS30ANTs as target template. Brain tissue segmentation of cerebrospinal fluid (CSF), white-matter (WM) and gray-matter (GM) was performed on the brain-extracted T1w using fast (10) (FSL 6.0.3:b862cdd5, RRID:SCR_002823). Brain surfaces were reconstructed using recon-all (11) (FreeSurfer 6.0.0, RRID:SCR_001847), and the brain mask estimated previously was refined with a custom variation of the method to reconcile ANTs-derived and FreeSurfer-derived segmentations of the cortical gray-matter of Mindboggle (12) (RRID:SCR_002438). Volume-based spatial normalization to two standard spaces (MNI152NLin6Asym, MNI152NLin2009cAsym) was performed through nonlinear registration with antsRegistration (ANTs 2.3.3), using brain-extracted versions of both T1w reference and the T1w template. The following templates were selected for spatial normalization: FSL\u2019s MNI ICBM 152 non-linear 6th Generation Asymmetric Average Brain Stereotaxic Registration Model (13) [RRID:SCR_002823; TemplateFlow ID: MNI152NLin6Asym], ICBM 152 Nonlinear Asymmetrical template version 2009c [(14) RRID:SCR_008796; TemplateFlow ID: MNI152NLin2009cAsym].

#### Functional data preprocessing

For each of the 1 BOLD runs found per subject (across all tasks and sessions), the following preprocessing was performed. First, a reference volume and its skull-stripped version were generated using a custom methodology of fMRIPrep. Head-motion parameters with respect to the BOLD reference (transformation matrices, and six corresponding rotation and translation parameters) are estimated before any spatiotemporal filtering using mcflirt (15) (FSL 6.0.3:b862cdd5). The estimated field map was then aligned with rigid-registration to the target EPI (echo-planar imaging) reference run. The field coefficients were mapped on to the reference EPI using the transform. BOLD runs were slice-time corrected to 0.972s (0.5 of slice acquisition range 0s-1.94s) using 3dTshift from AFNI (16) (RRID:SCR_005927). The BOLD reference was then co-registered to the T1w reference using bbregister (FreeSurfer) which implements boundary-based registration (17). Co-registration was configured with six degrees of freedom. Several confounding time-series were calculated based on the preprocessed BOLD: framewise displacement (FD), DVARS and three region-wise global signals. Global signal was extracted within the whole-brain masks. Additionally, a set of physiological regressors were extracted to allow for component-based noise correction (18) (CompCor). Principal components are estimated after high-pass filtering the preprocessed BOLD time-series (using a discrete cosine filter with 128s cut-off) for the anatomical (aCompCor). For aCompCor, three probabilistic masks (CSF, WM, and combined CSF+WM) are generated in anatomical space. The implementation differs from that of Behzadi et al. in that instead of eroding the masks by 2 pixels on BOLD space, the aCompCor masks are subtracted from a mask of pixels that likely contain a volume fraction of GM. This mask is obtained by dilating a GM mask extracted from the FreeSurfer aseg segmentation, and it ensures components are not extracted from voxels containing a minimal fraction of GM. Finally, these masks are resampled into BOLD space and binarized by thresholding at 0.99 (as in the original implementation). Components are also calculated separately within the WM and CSF masks. For each CompCor decomposition, the k components with the largest singular values are retained, such that the retained components time series are sufficient to explain 50 percent of variance across the nuisance mask. The remaining components are dropped from consideration. The BOLD time-series were resampled into standard space, generating a preprocessed BOLD run in MNI152NLin6Asym space. First, a reference volume and its skull-stripped version were generated using a custom methodology of fMRIPrep. Gridded (volumetric) resamplings were performed using antsApplyTransforms (ANTs), configured with Lanczos interpolation to minimize the smoothing effects of other kernels (19). Finally, the aCompcor, cosine (highpass filtering), six head-motion and global signal regressors were regressed out of each subjects MNI-space voxel level images.

#### Computing individual-level functional connectivity matrices

To characterize the functional profile of CUD for each individual, we used previously validated Schaefer 400 cortical (20) and Tian 32-Scale II subcortical (21) atlases (**Fig. 1A**) to extract regional time series by taking the average of all voxels belonging to a given region. We then calculate the pairwise Pearson correlation between each of the 432 regions, to generate 93,096 edge functional connectivity matrix. We employed the Yeo 7-network parcellation (22) to assign each cortical ROIs to a corresponding functional network. Subcortical regions were classified according to their broad-scale anatomy (21).

### Section 4. Methods

#### Network Based Statistic

The Network-Based Statistic (NBS) method tackles the challenge of multiple comparisons that arises in the context of whole brain connectome analyses. It accomplishes this by conducting statistical assessments at the level of interconnected components, which comprise groups of nodes linked together through a series of edges, in contrast to the conventional treatment of each individual edge in isolation. Specifically, at each edge (i.e., functional connectivity estimates between two regions), differences in functional connectivity between individuals with CUD and matched controls were assessed using an ANCOVA, adjusting for age, sex, and education, examining the main effect of the group. Using R-version-4.0.3, package NBR (R package version: 0.1.5) (23), the Network Based Statistic (NBS) was used to perform family-wise error-corrected (FWE) inference at the level of connected-components of edges showing a common effect, with significance assessed at *p_FWE_* < 0.05. The NBS procedure involves setting a primary component-forming threshold (τ), which is applied to both the observed data, and the permuted null data. The decision on where to set this threshold is arbitrary; a lower threshold can detect weaker differences over many edges, whereas a higher threshold tends to pinpoint stronger effects that might span fewer edges. We report results for τ < 0.05 here, and present results for and τ < 0.01, and τ < 0.001 in **Supplementary Table 2 and Supplementary Figure 1**. For both the observed and permuted null data, we noted the size (number of edges) in the connected components above this threshold. The size of the largest component from each permutation was employed to construct a null distribution, and a corrected-value for each observed component was estimated as the proportion of null component sizes that was larger than the observed value (24). To comprehensively delineate brain-wide alterations in functional connectivity, we present the results at three different scales: (1) the level of individual connections (i.e., where edges are either under- or over-connected, or hypo- versus hyperconnectivity, respectively); (2) the level of individual brain regions, to identify specific brain areas which had a high number of significant connections (**Fig. 1B, 1D**); and (3) the level of large-scale functional brain networks, analyzed both within- and between-network. Here, we examined both as proportion of implicated edges (e.g., upper triangle **Fig. 1C, 1E**) and as proportions normalized by the size of the network (e.g., lower triangle **Fig. 1C, 1E**). To determine whether the observed functional connectivity alterations showed any network-specificity, we calculated the proportion of significant edges that fell within each brain networks (e.g., **Fig. 1C**; upper triangle). Different brain networks have intrinsic differences in their size (number of regions), therefore we present both raw proportions and proportions normalized by the total number of possible network connections between each pair of networks (e.g. **Fig. 1C** lower triangle of matrix); the former identifies preferential involvement of a given network in an absolute sense while the latter accounts for differences in network size (i.e., the tendency for larger networks to be more likely to be implicated in a given NBS network).

#### Neuromaps

Receptor density data were obtained from Neuromaps (25). Neuromaps is an open source a toolbox for accessing and analyzing structural and functional brain maps, combined from open-access data to compare brain maps. Each group-level parametric PET image was parcellated into 432 regions using the same atlases as the functional MRI data, and these regional values were z-scored within each map. To quantify the relationship between the various receptor distributions and CUD-related functional alterations, we first computed the degree of the detected NBS network (number of significant edges connecting each region). We then performed Spearman’s correlation between regional degree and each receptor expression, using ‘spin tests’ for non-parametric inference that accounts for spatial autocorrelation (10,000 permutations; (26)). Subcortical regions were random shuffled within hemisphere at each permutation (27, 28). Specifically, to evaluate receptor density map comparison significance, a corresponding set of null models was generated by computing 10,000 null correlations between permuted degree from the detected NBS network (see Methods) and observed receptor densities. The p-value for each correlation’s significance is defined as the proportion of null models with correlation values greater the original (observed) value. Binding potential refers to the ratio at which a radioligand binds to a specific receptor within the brain compared to its nonspecific binding.

**Supplementary Table 1:**
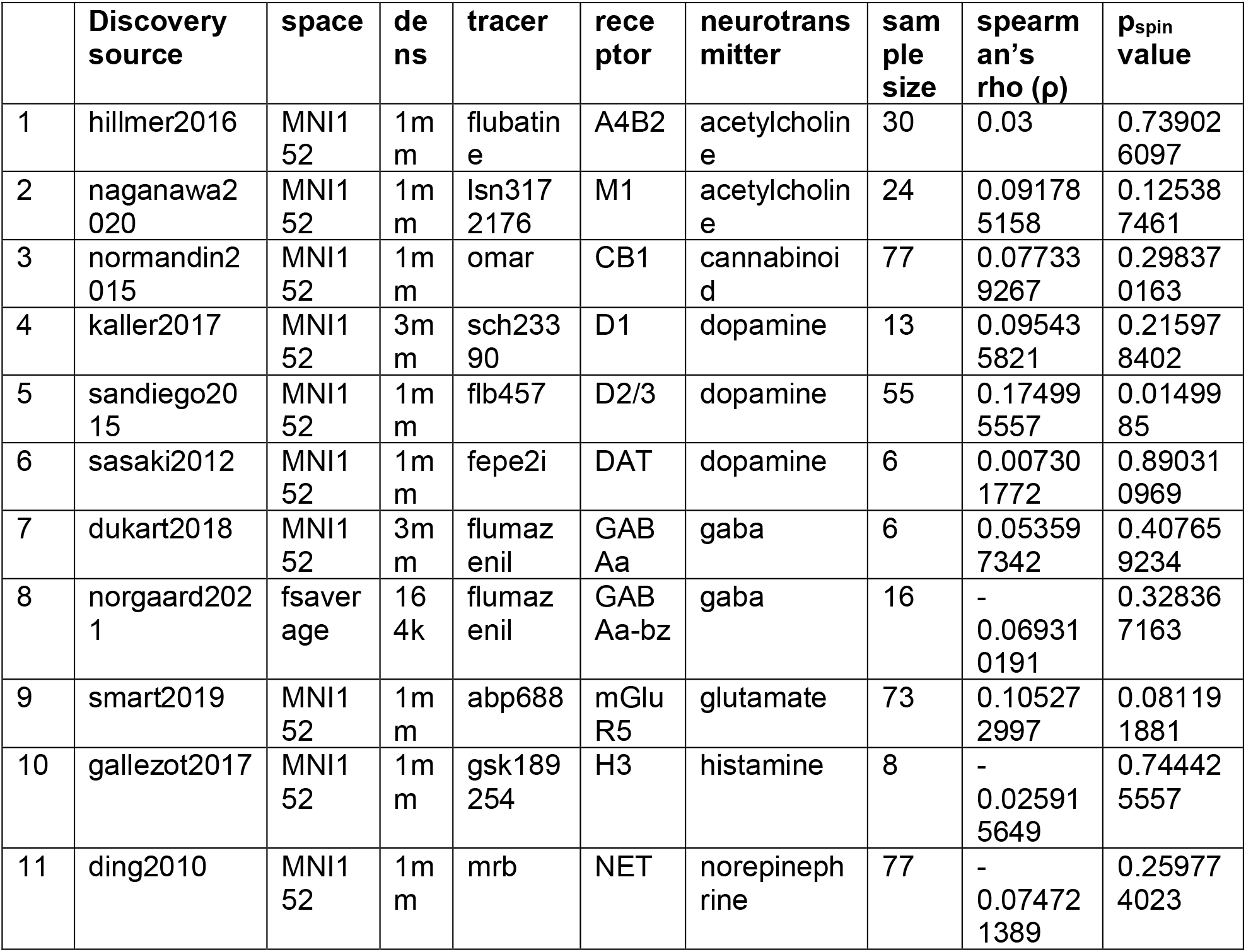
Results using alternate component forming threshold (τ=0.01).

**Supplementary Table 2:**
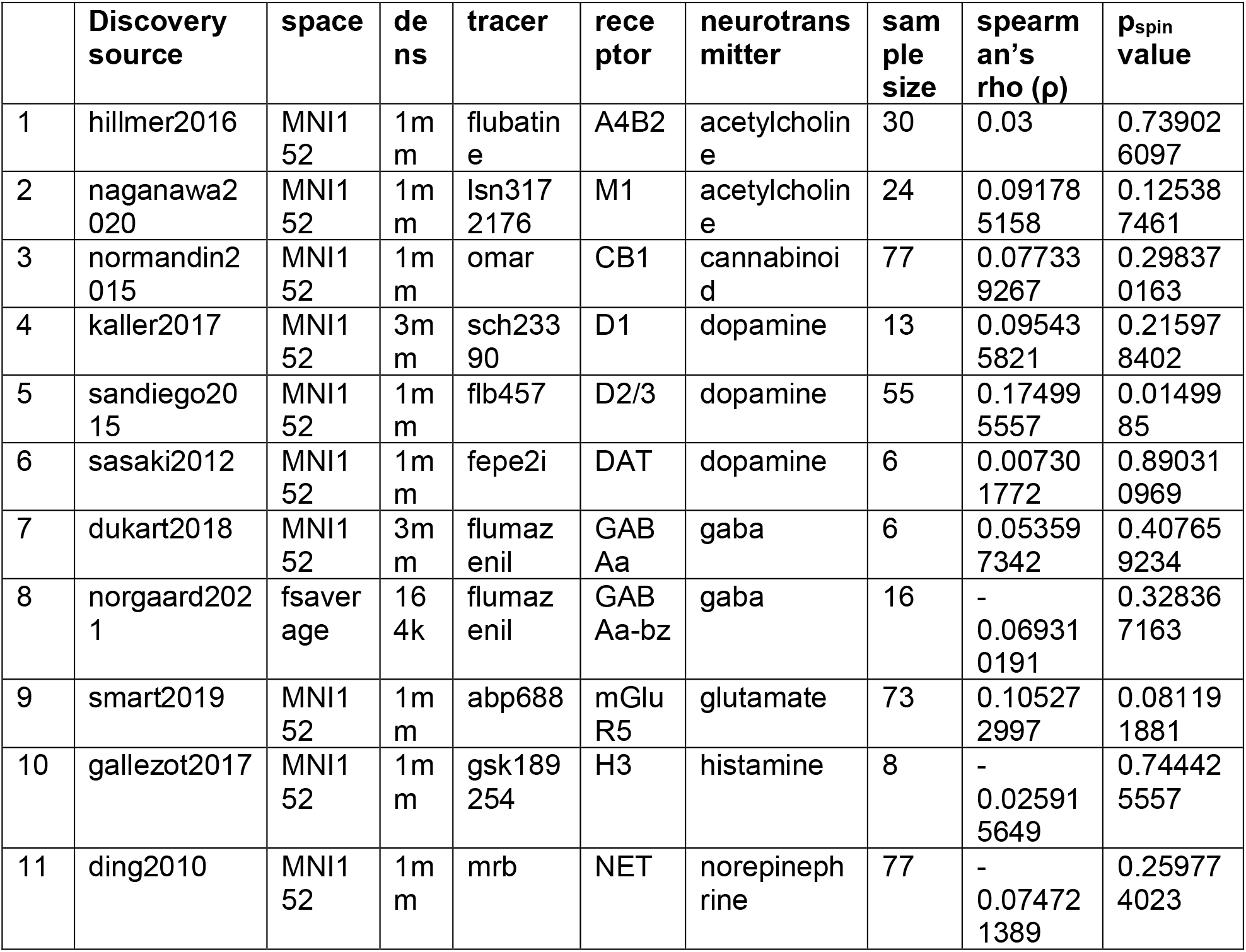
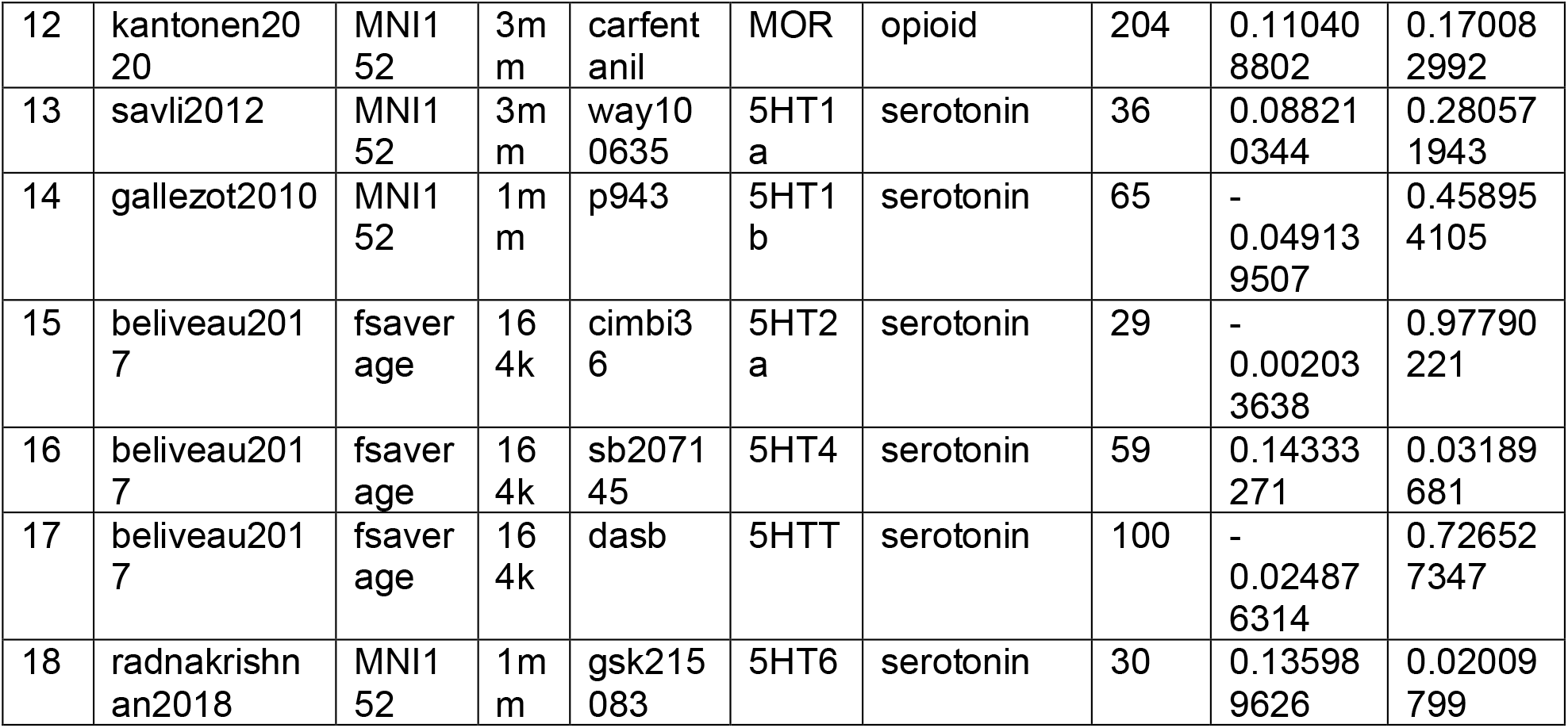
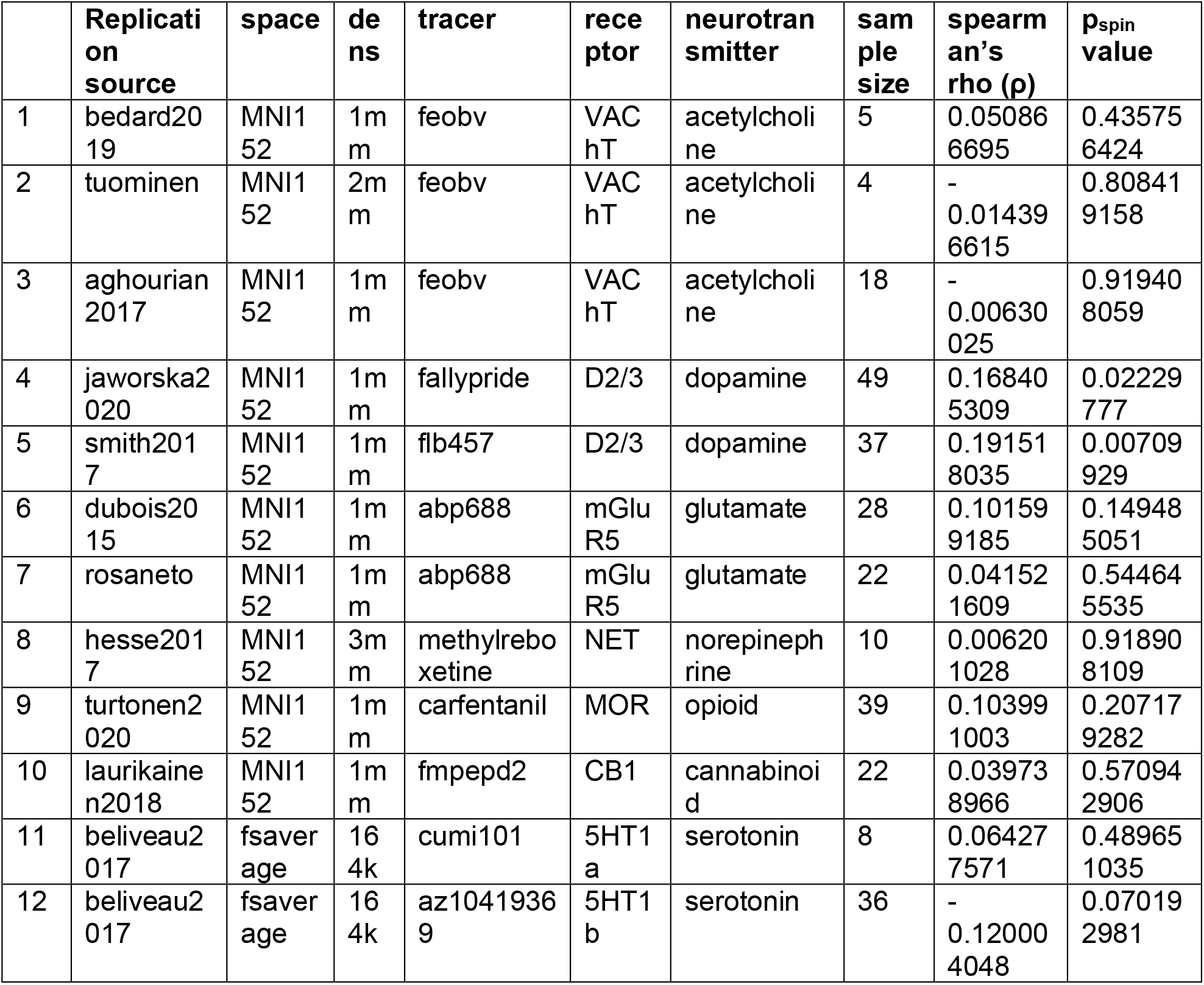
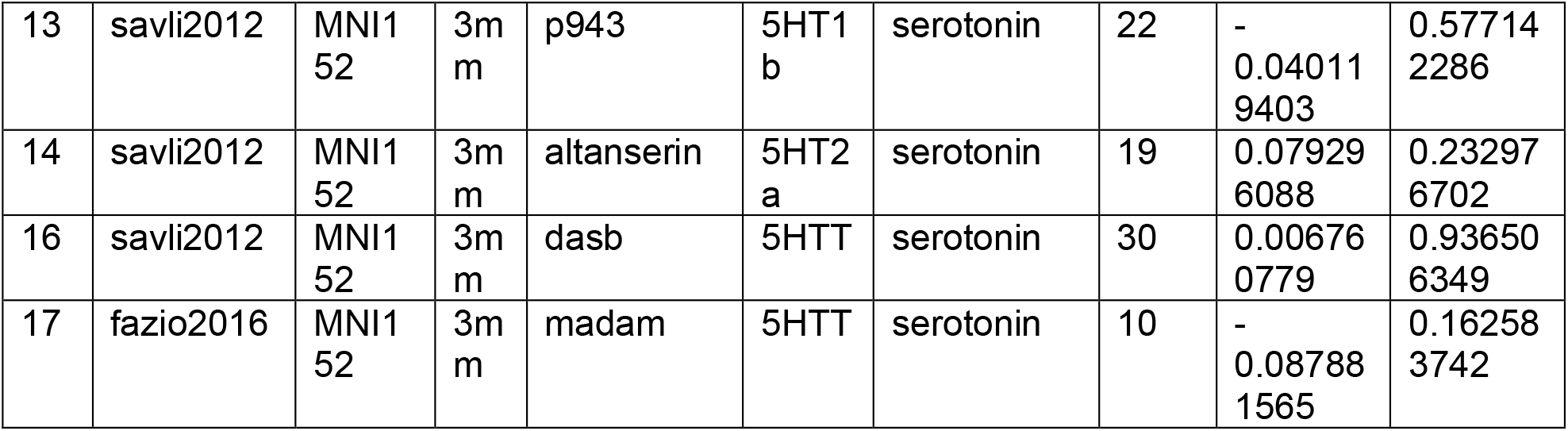
Neurotransmitter receptor density spearman’s ρ correlations after 10k spin test and random shuffling per hemisphere for subcortical regions, and p_spin_-values (uncorrected) for both 1) non-duplicate discovery set of receptor maps and 2) replication receptor maps. Raclopride D_2/3_ tracer was excluded due to inconsistent binding as previously reported using neuromaps (32).

**Supplementary Figure 1:**
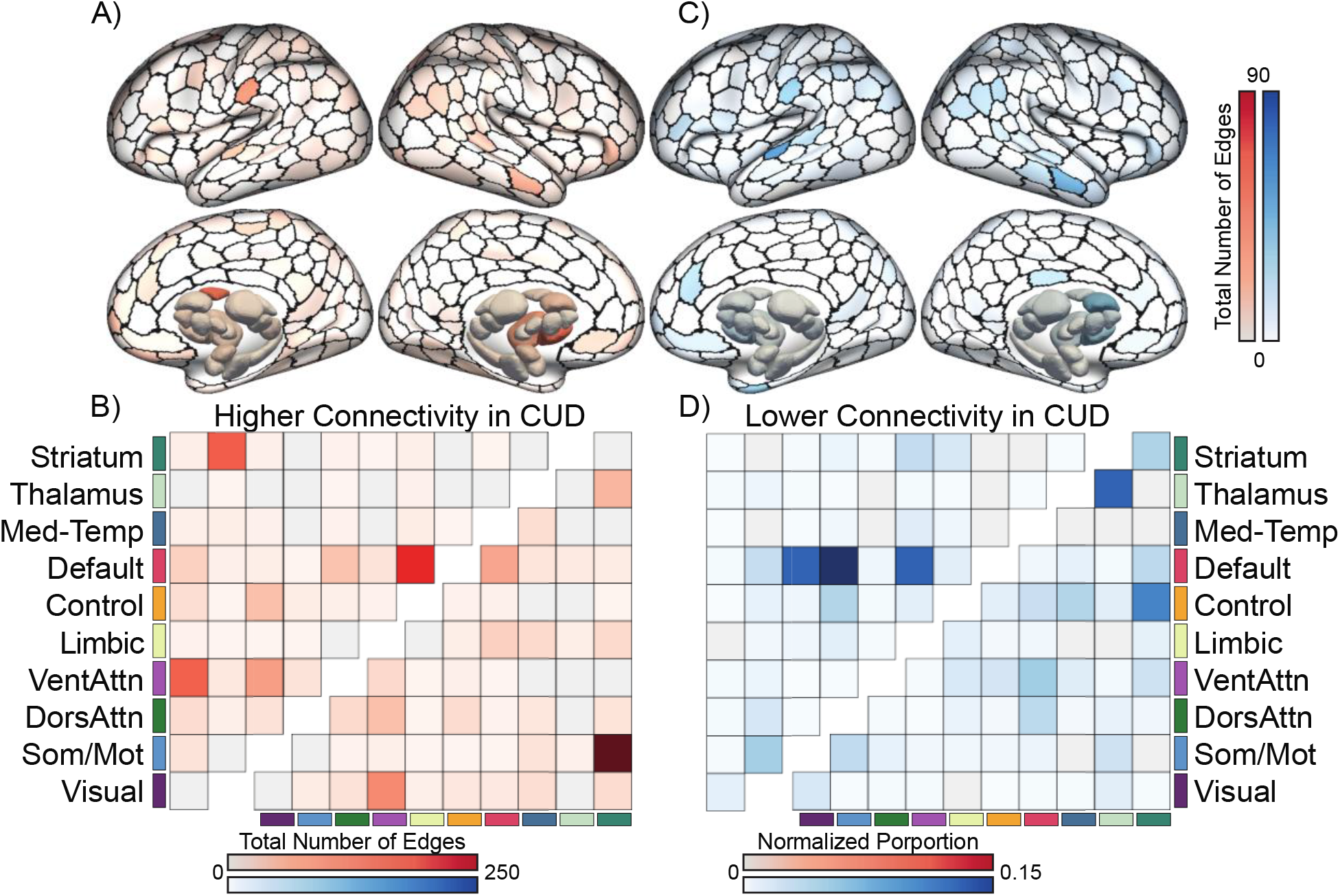
Results using alternate component forming threshold *τ =* 0.01. No significant component was detected at τ < 0.001, suggesting that the detected effect is disperse and spatially widespread.

**Supplementary Figure 2:**
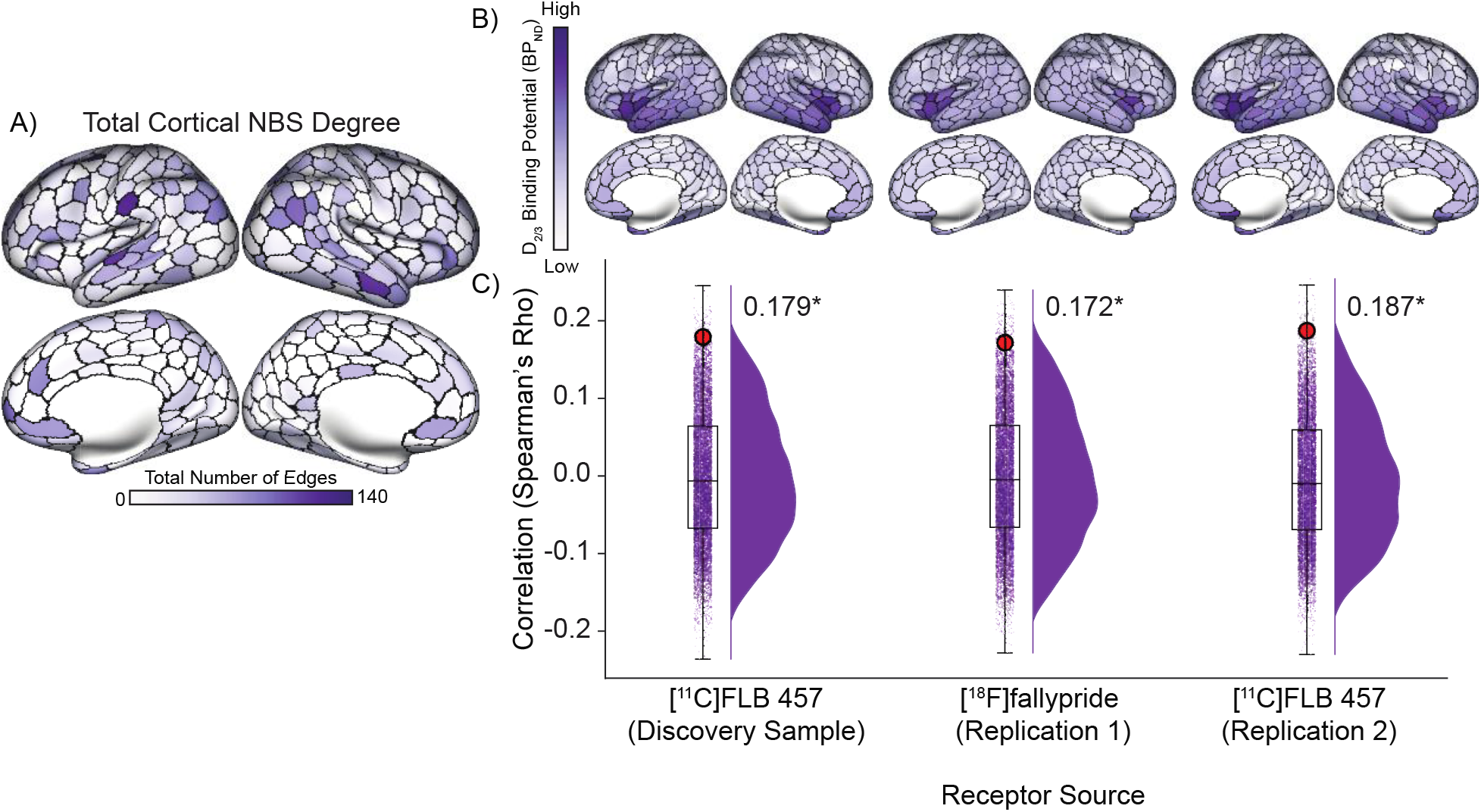
Results remain significant when subcortical regions are excluded. Regional dysconnectivity was significantly correlated with D_2/3_ receptors in the absence of subcortical only associations. Color scale for PET maps normalized between -1.0 to 1.0 for cortex and subcortex separately. (Sandiego et al., 2015. (29) Discovery Sample: ρ=0.179; p_spin_=0.018); (Jaworska et al., 2020. (30) Replication 1: ρ=0.172; p_spin_=0.029); (Smith et al., 2017 (31): Replication 2: ρ=0.187; p_spin_=0.013)

**Supplementary Figure 3:**
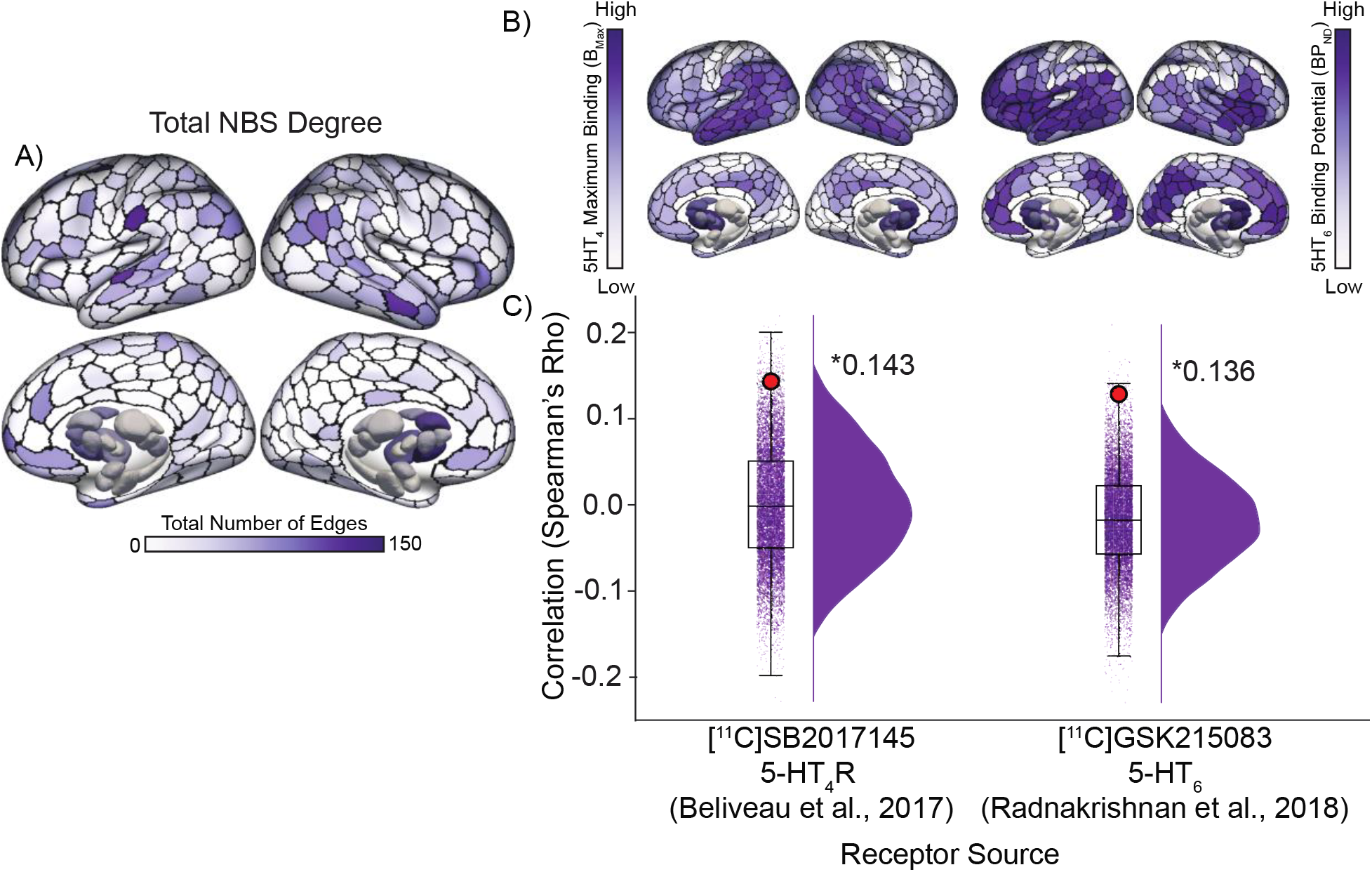
NBS total degree dysfunction associations with serotonin receptors. Regional dysconnectivity was additionally significantly correlated with 5HT_4_ (ρ=0.143; p_spin_=0.032), and 5HT_6_ (ρ=0.136; p_spin_=0.020) serotonin receptors, however, replications were not available. Color scale for PET maps normalized between -1.0 to 1.0 for cortex and subcortex separately. (5-HT_4_-R: Beliveau et al., 2017 (33)) (5-HT_6_: Radhakrishnan et al., 2018 (34)). 5HT_4_: B_Max_ was converted from BP_ND_ using autoradiography densities.

